# Identification and verification of the potential effect of therapeutic miRNA-mRNA pairs on ferroptosis in small cell lung cancer by bioinformatics analysis

**DOI:** 10.1101/2021.06.28.450166

**Authors:** Yuefeng Wu

## Abstract

Owing to the high mortality rate of small cell lung cancer (SCLC), it is essential to determine a novel therapeutic approach for treating patients with SCLC. miRNA is a type of non-coding RNA that plays a role in translational control. By applying this identity, it is applicable for treating patients using miRNA and nanotechnology. Ferroptosis is a newly discovered type of programmed cell death. Accumulated evidence suggests the possibility of using ferroptosis in treating patients with SCLC. Thus, identifying potential therapeutic miRNA-mRNA pairs that have an impact on ferroptosis will be valuable. In the in silico analysis, several Gene Expression Omnibus datasets were analyzed. The results were verified using other data. Here, we report that the miR-30 family and KIF11 pair have a crucial influence on SCLC cells, which may also affect ferroptosis.

## 1. Introduction

Small cell lung cancer (SCLC) is a notable global disease with high mortality and a 5-year survival rate of less than 7% in the past decades (Byers and Rudin 2015). SCLC is characterized by breathing difficulties and a unique type of pain due to the rapid progression of cancer cells and some paraneoplastic syndromes. Current chemotherapy medication, such as etoposide or irinotecan, cannot be satisfactorily used to treat patients. (van Meerbeeck, Fennell, and De Ruysscher 2011) The development of advanced precision medicine technology has made it possible to treat patients based on specific biomarkers. Both the technology situation and the disease yield a bioinformatics analysis for SCLC data. Therefore, potential drug targets or biomarkers are required.

Meanwhile, SCLC accounts for only approximately 14% of all lung cancers (Byers and Rudin 2015), with the remaining being non-small cell lung cancer (NSCLC). However, compared with SCLC therapy, there have been some advances in targeted therapies for NSCLC. Several miRNA-mRNA pairs have been identified and validated using bioinformatics considering the insufficient data on NSCLC.

However, the lack of research material (e.g., DNA, RNA, tissues, etc.) has limited translational research on SCLC, which is mainly caused by the rapid disease progression and other factors (Byers and Rudin 2015). Even in recent years, although there are some SCLC databases, bioinformatics analysis on SCLC is still limited; hence, such bioinformatics analysis will be valuable.

MicroRNAs can regulate gene expression (Lee and Dutta 2009). Abnormal gene expression has been recognized as the cause of a wide range of cancers. Moreover, miRNA-based anti-cancer therapy, which can be used along with chemotherapy, has been discussed over the years. Therefore, it is essential to identify suitable hub genes and miRNAs. Hence, it would be useful to determine correlations between these miRNAs and mRNAs (Hayes, Peruzzi, and Lawler 2014).

In this study, we aim to identify and verify the potential therapeutic miRNA-mRNA pairs for SCLC by bioinformatics analysis using datasets obtained from the Gene Expression Omnibus (GEO) database.

First, hub genes are identified by analyzing the data from the RNA-seq. Pathway enrichment is performed as normal. Differentially expressed miRNAs are also identified. These miRNAs are enriched in terms of their functions and target genes. KIF11 is shown in both the hub gene set and the differentially expressed miRNA set. DNA methylation also influences DNA transcription; therefore, the KIF11 methylation status has also been reported. Overall, we report that the miR-30 family plays an essential role in SCLC by interacting with KIF11.

Moreover, unsupervised cluster analysis shows the potential effect of KIF11 on ferroptosis. Ferroptosis is a type of programmed cell death that results from an irreparable lipid peroxidation chain reaction within cellular membranes fueled by radical formation (Tang et al. 2020). It has drawn much attention in recent years because it provides a novel therapeutic approach. This correlation has made the discovery of our miRNA-mRNA pairs more valuable.

## 2. Materials and Methods

### 2.1 Microarray data

Datasets from the GEO database (Barrett et al. 2013) were obtained using the following criteria: 1) SCLC, 2) patients, 3) miRNA, or 4) mRNA. GSE40275 (Kastner et al. 2012) contains gene expression datasets from samples derived from normal lungs and patients with SCLC and NSCLC. A total of 43 normal samples were included in the wild type group and 24 samples were distributed into the SCLC groups. These two groups were used for differentially expressed gene analysis. GSE19945 contains miRNA profiling files with 35 samples of SCLC and eight samples of normal lung tissue obtained and used for further investigation. GSE50412 (Saito et al. 2016) is a methylation profile containing 28 fresh frozen samples and 13 non-cancerous lung tissues. The corresponding probe annotations were downloaded for further analysis. SCLC cell line datasets were downloaded from http://sclccelllines.cancer.gov. Further sorting and searching were completed by the navigation of the SCLC project website.

### 2.2 Differentially expressed gene detection and enrichment analysis

The data from the MINIML file were extracted, normalized, and processed by log2 transformation using the preprocessCore package in R software (version 3.4.1). Probes were converted to gene symbols according to the platform annotation information of the normalized data. Probes with more than one gene were eliminated, and the average value was calculated for genes corresponding to more than one probe. The batch effect was removed using the removeBatchEffect function of the Limma package in R.

The Limma package (version: 3.40.2) was used to study the differential expression of mRNAs. “Adjusted P < 0.05 and Log (Fold Change) >1 or Log (Fold Change)< −1” were defined as the thresholds for the screening of differential expression of mRNAs. The ClusterProfiler package (version: 3.18.0) in R was used to analyze the GO function of potential targets and enrich the Kyoto Encyclopedia of Genes and Genomes (KEGG) pathway (Yu et al. 2012).

The top 1000 differentially expressed genes (DEGs) were identified and searched in the STRING (https://string-db.org/) database for protein-protein interaction (PPI). Cytoscape was used to construct a PPI network. Cytoscape and CytoHubba with the MCC algorithm were used to analyze the hub genes and visualize the molecular interaction networks.

### 2.3 Unsupervised cluster based on hub genes and ferroptosis

The R software package ConsensusClusterPlus (v1.54.0) was applied for consistency analysis, the maximum number of clusters is 6, and 80% of the total sample is drawn 100 times, clusterAlg = “hc,” innerLinkage=‘ward.D2’. The R software package pheatmap (v1.0.12) was used to cluster the heatmaps. The gene expression heatmap retains genes with SD > 0.1(Yi et al. 2020).

### 2.4 DEM detection and enrichment analysis

The miRNA data were processed using GEO2R. The groups were distributed as described above. The results obtained from GEO2R were visualized using ggplot2. The miRNA enrichment analysis and workflow management system miEAA was used for DEM enrichment and target gene prediction (Kern et al. 2020). The top 100 DEMs were selected, and their names were translated as per platform instructions. Then, an over-representation analysis was performed. Gene ontology (miRWalk), pathways (miRWalk) (Sticht et al. 2018) and target genes (miRTarbase) (Hsu et al. 2011) were chosen as parameters for analysis. The significance level was set as 0.05, whereas the threshold level was set as 2.

## 3. Results

### 3.1 DEG detection and enrichment analysis

The analysis successfully identified 3739 DEGs, among which 1708 genes were downregulated and 2031 genes were upregulated. The 1000 genes with the lowest p-values were selected for PPI network construction. With the help of MCC, the top 14 hub genes were obtained, which all ranked first in the list (Fig. 1). These are CCNB1, CDK1, NCAPG, KIF11, BUB1B, NUSAP1, TTK, CCNB2, MELK, TOP2A, CCNA2, PBK, BUB1, and KIF20A. KEGG pathway enrichment revealed that the pathways related to RNA regulation, DNA replication, cell cycle, and p53 were upregulated. In contrast, adhesion ability, TNF signaling pathway, and PI3K-Akt signaling pathway were downregulated. The GO term enrichment showed that the terms enriched in cell cycle, DNA conformational change, nuclear division, and other mitosis-related terms were upregulated. Meanwhile, the terms related to extracellular matrix and structure organization, vasculature development, cell adhesion, and immune system processes were downregulated. The results provided an overview of the SCLC landscape (Fig. 2).

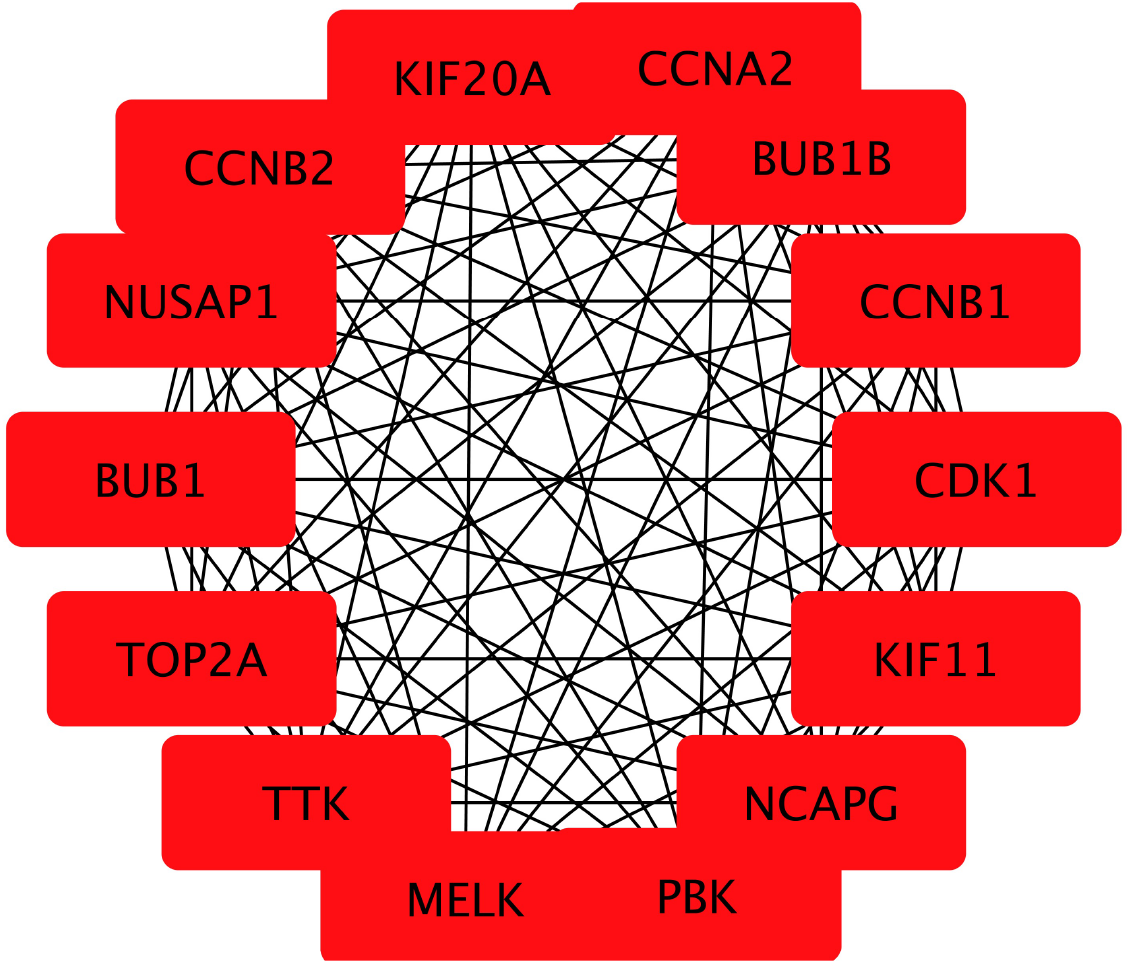

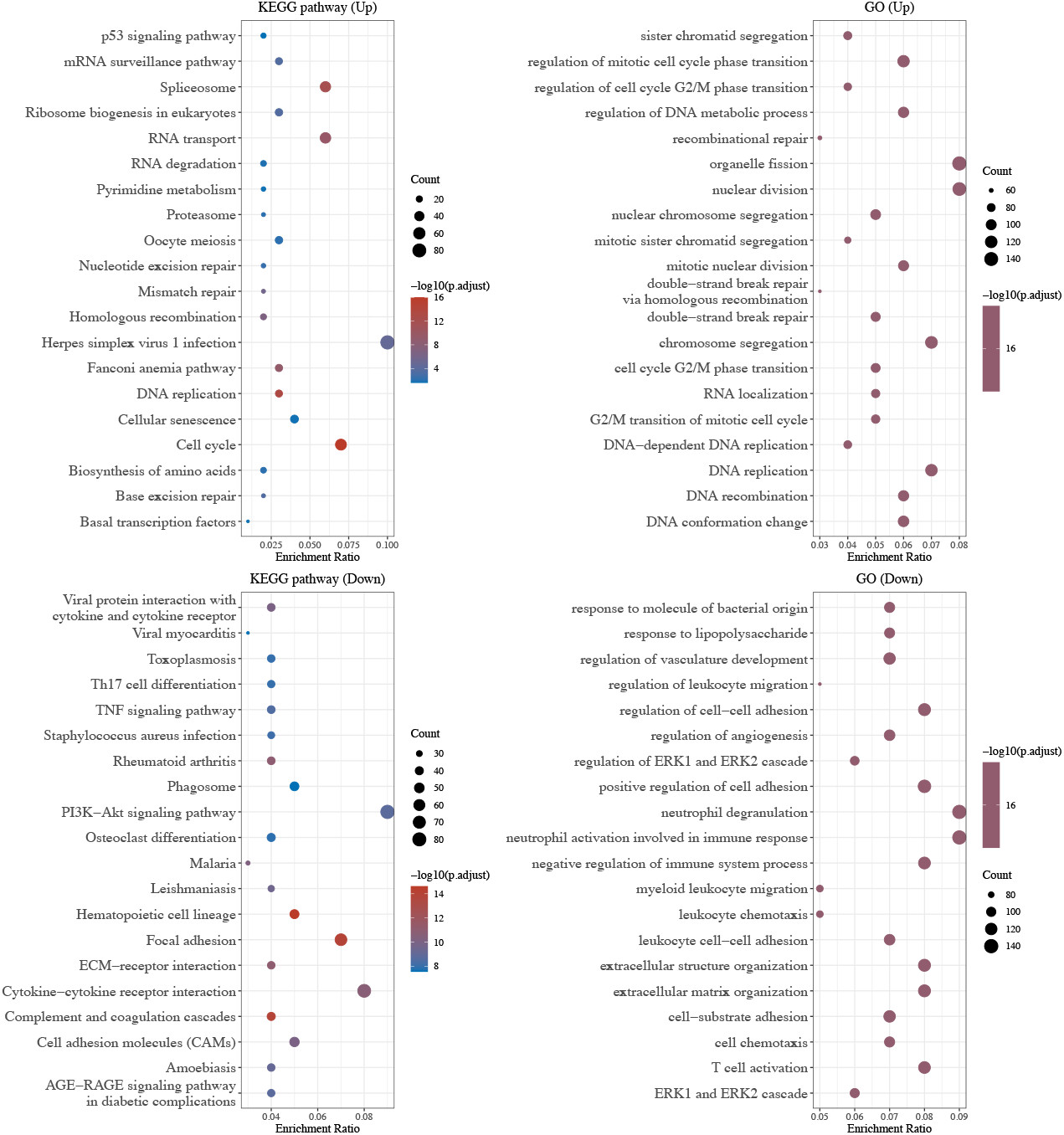

### 3.2 Unsupervised cluster based on hub genes and ferroptosis

There are three different groups (Fig. 3) sorted based on the ferroptosis gene list. However, one group has fewer than two elements, which may be coincidental. Some types of SCLC cells are sensitive to ferroptosis and another type to contrast (Bebber et al. 2021). Therefore, it is reasonable to combine the three groups into two groups. The two groups were obtained by sorting using the hub gene list and following the previous logic (Fig. 4). By comparing the elements from the corresponding groups, most were found to be identical. The PPI network of hub genes and ferroptosis genes were obtained; KIF11 interacts with GPX4. Furthermore, the difference in ferroptosis gene expression between the unsupervised cluster group and the hub gene list showed significant differences (Fig. 5) (Liu et al. 2020).

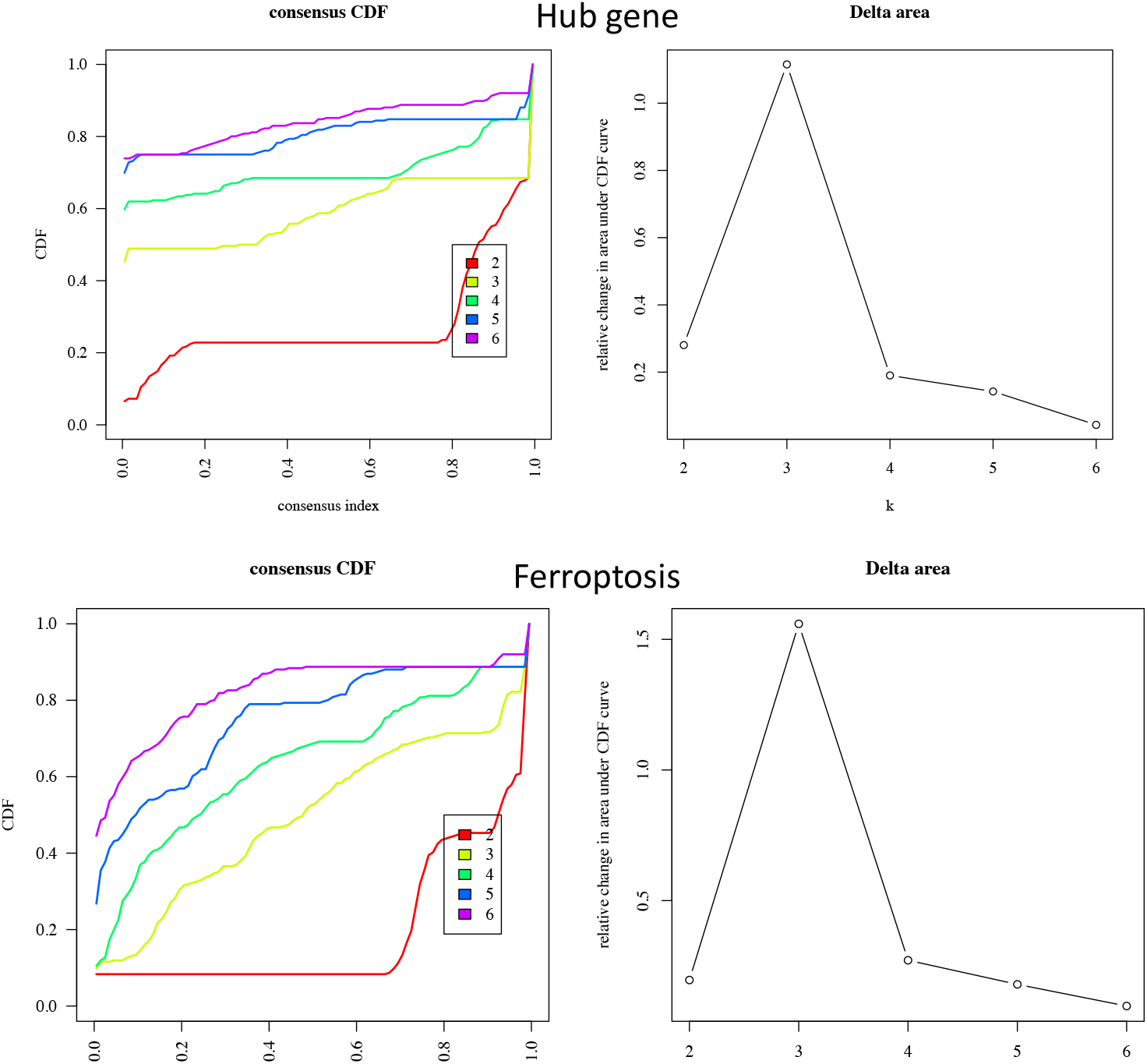

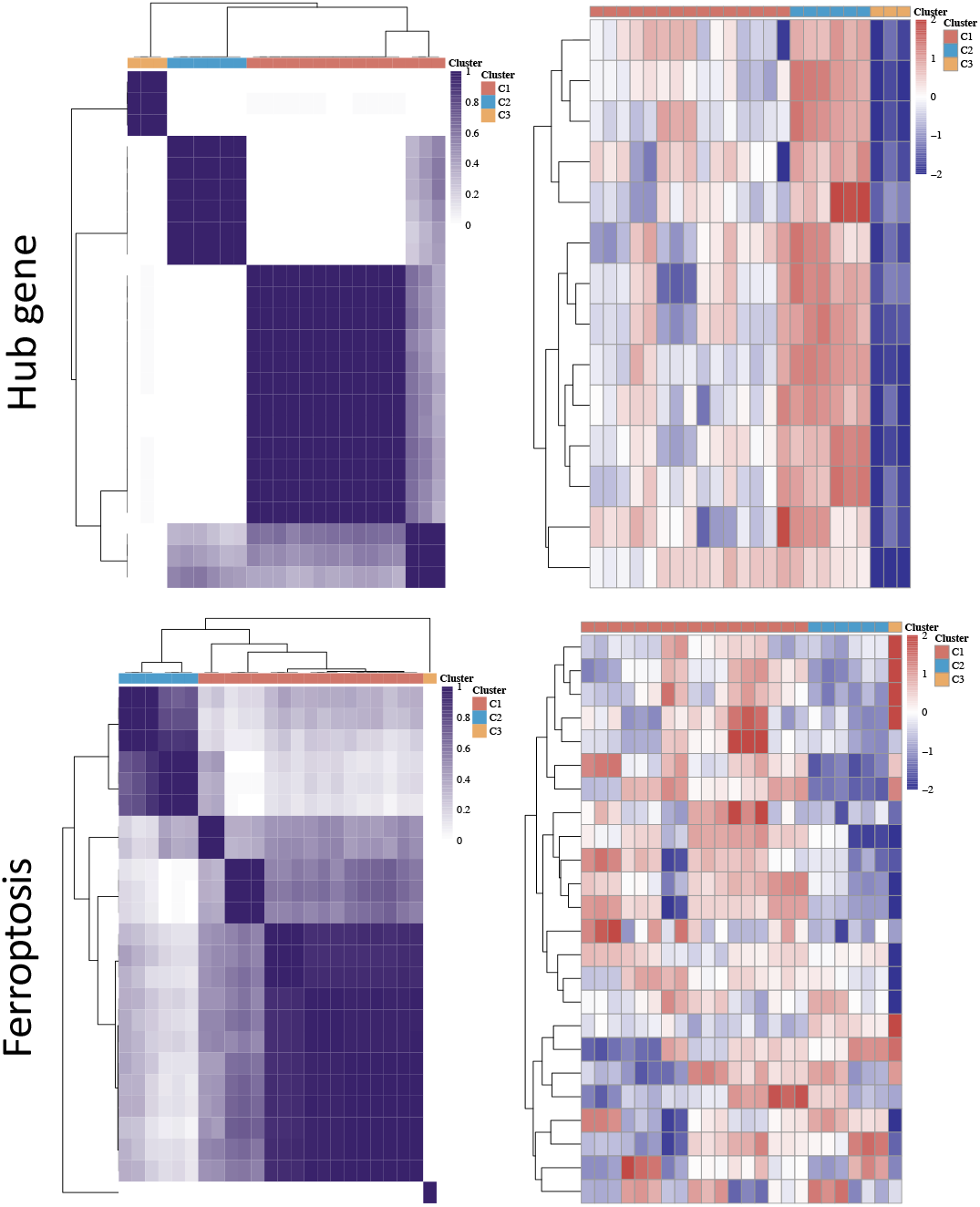

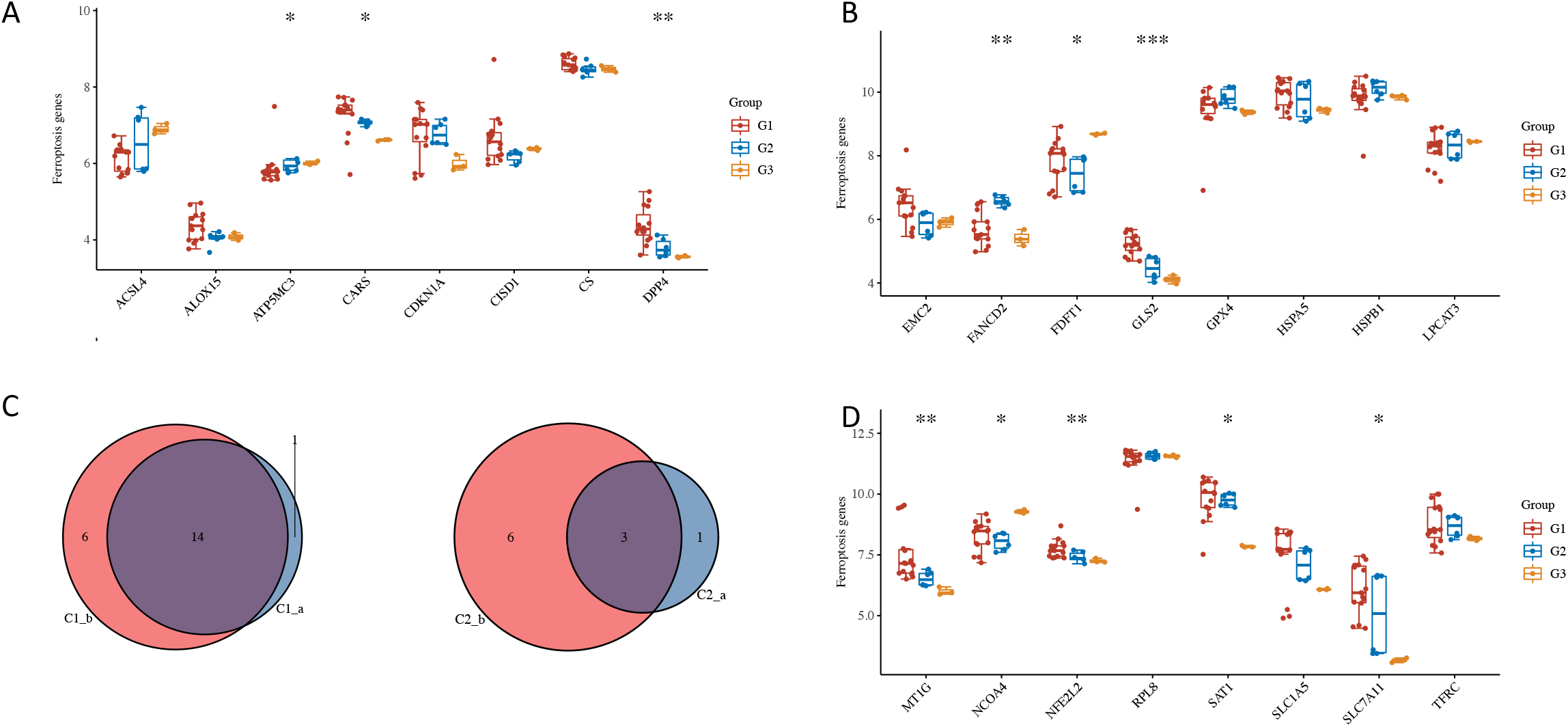

### 3.3 DEM detection and enrichment analysis

Differentially expressed miRNAs were successfully obtained using GEO2R (Fig. 6). The table contains p-values, IDs, sequences, etc. These data were visualized using ggplot2. The top 100 miRNAs with the lowest p-values were selected and uploaded to miEAA (Fig. 7). These miRNAs were mainly enriched in terms of PD-L1 expression, PD-1 checkpoint, VEGF signaling pathway, and some hormone-related pathways. Target gene prediction successfully obtained 1271 target genes. In fact, two of these genes—CCNA2 and KIF11—also appeared in the hub genes. The miR-30 family is believed to influence KIF11 expression (Fig. 8).

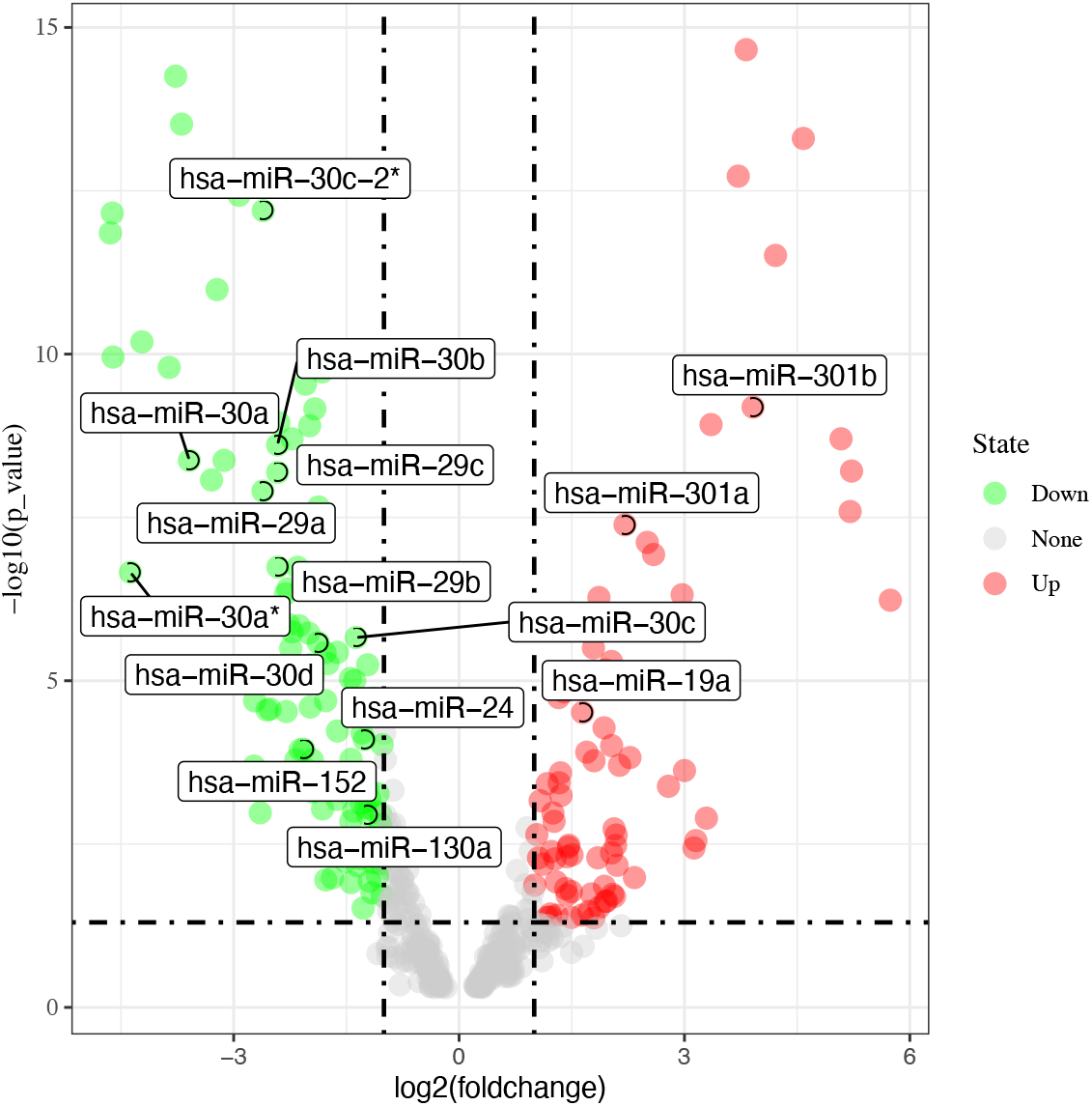

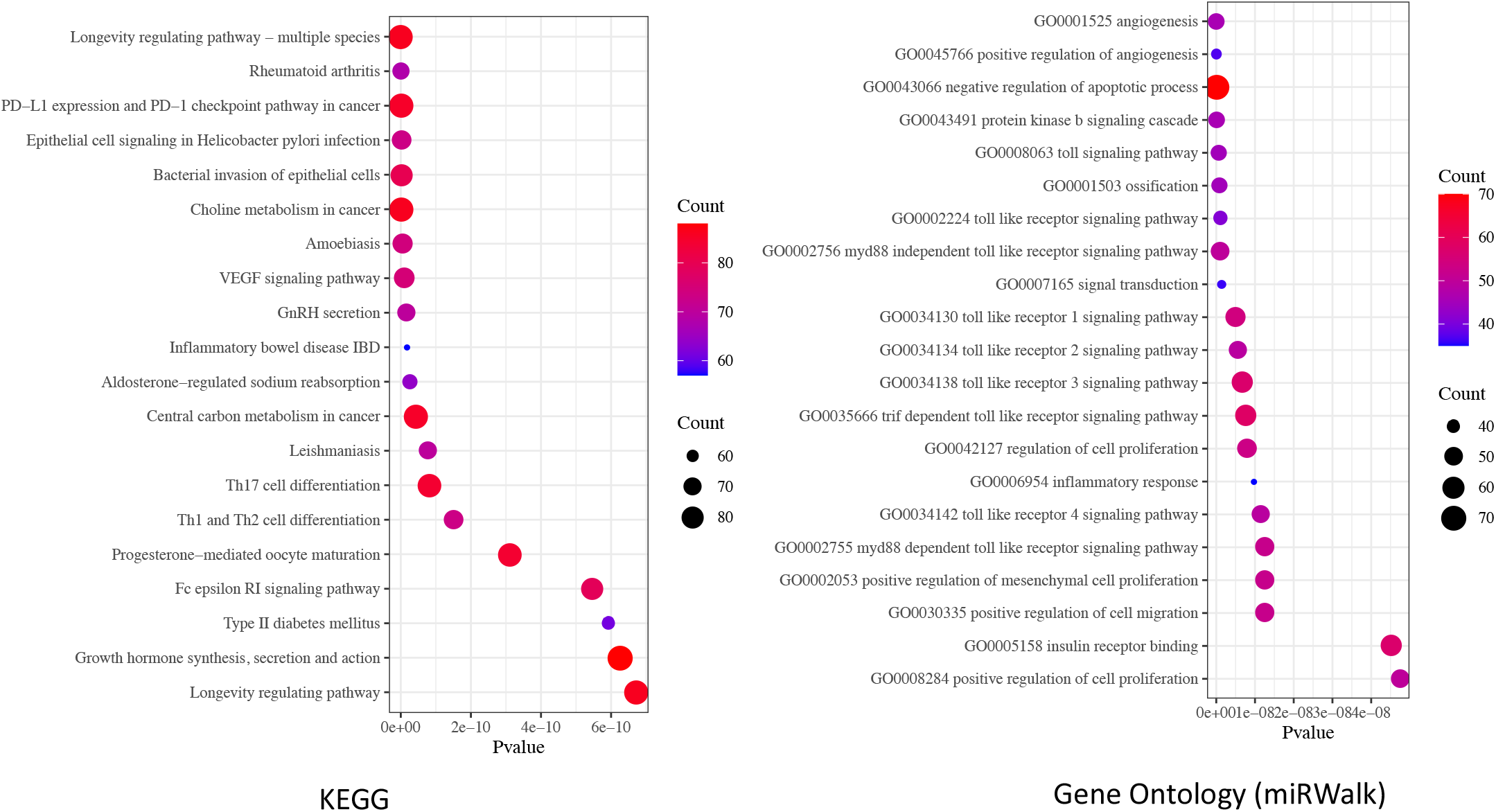

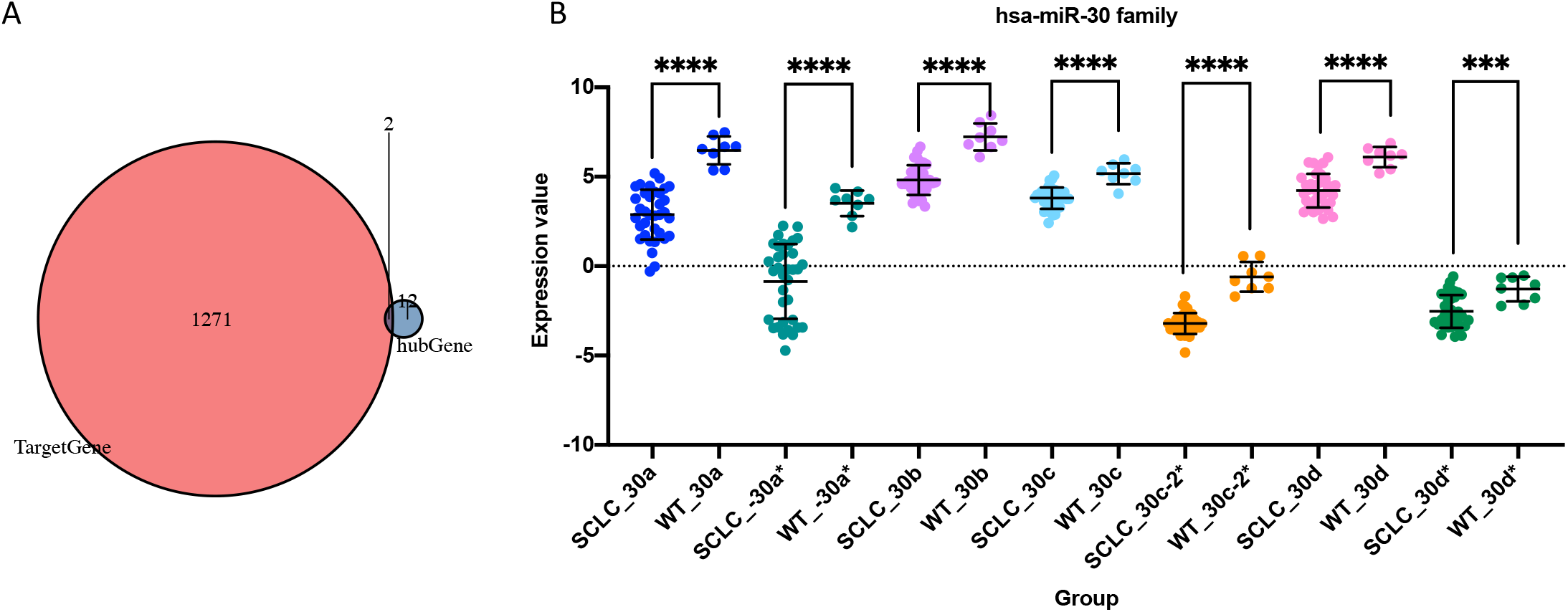

### 3.4 KIF11 methylation analysis

The dataset containing SCLC methylation profiling was imported into GEO2R and run. The corresponding platform annotation list was used to obtain the probe ID for KIF11; cg25494789 and cg16340918 are two probes with a KIF11 tag. Their methylation status in the SCLC and normal groups was compared using a t-test (Fig. 9). No statistical significance was found for cg25494789. The p-value was 0.0271 for the comparison of cg16340918. Although there was some statistical significance, it was not as significant as the difference in the miR-30 family.

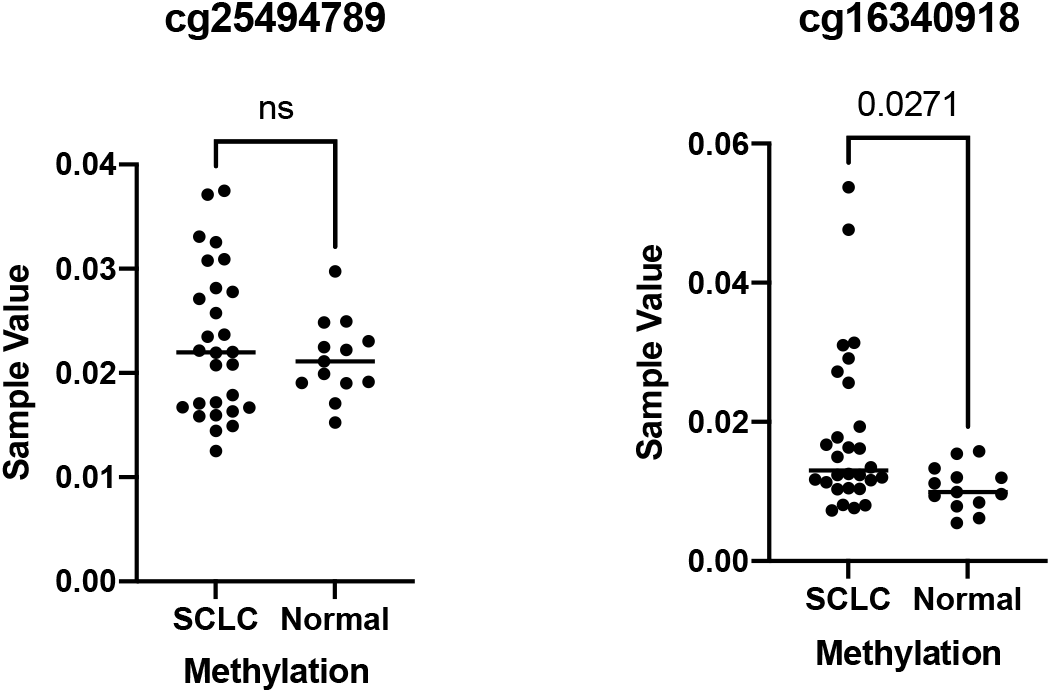

## 4. Discussion

Owing to the high malignancy characteristics of SCLC, it is essential to identify and verify the biomarkers for SCLC. Although miRNA cannot code for any protein, it can bind to the mRNA to either slow down translation or degrade the mRNA. Abnormal expression of miRNA has been found to be related to various cancers. It is also a promising tool for cancer therapy (Mishra, Yadav, and Rani 2016).

In this in silico analysis, the DEGs were first identified, and some crucial pathways were presented. P53 signaling plays an important role in the coordination of cellular responses to different types of stress, such as DNA damage and hypoxia. Activation of p53-regulated genes leads to cellular senescence, cell cycle arrest, or apoptosis.

The TNF signaling pathway is associated with tumor necrosis. The abnormal alteration of this pathway affects immune and non-immune cell types, including macrophages, T cells, mast cells, granulocytes, natural killer (NK) cells, fibroblasts, neurons, keratinocytes, and smooth muscle cells as an immune response to pathogenic stimuli (Blaser et al. 2016). The PI3K-Akt signaling pathway is activated when PI3K phosphorylates Akt, thereby activating it. Once activated, Akt controls a number of downstream cellular processes, including apoptosis, protein synthesis, metabolism, and cell cycle, by phosphorylating a range of substrates (Martini et al. 2014). The AGE-RAGE signaling pathway influences NAPDH and enhances oxidative stress, which activates the NF-κB signaling pathway and further stimulates the production of cytokines and growth factors, thereby damaging cells and tissues (Waghela et al. 2021). These findings are consistent with the findings of GO enrichment.

Most hub genes that have been identified are related to the cell cycle. KIF11 encodes a motor protein belonging to the kinesin-like protein family. Members of this protein family are known to be involved in various kinds of spindle dynamics. The function of this gene product includes chromosome positioning, centrosome separation, and establishment of a bipolar spindle during cell mitosis (Asbaghi et al. 2017).

The miRNA enrichment results pertain to T cell differentiation and certain immune checkpoint pathways. Target gene enrichment revealed that the miR-30 family has a crucial impact on KIF11, which is also an obtained hub gene; this is important information as the development of nanotechnology can facilitate delivery of miRNA into cells to control the hub gene. It has been reported that the miR-30 family plays a crucial regulatory role in the development of tissues and organs and the pathogenesis of clinical diseases, indicating that it may be a promising regulator in development of the disease (Mao et al. 2018).

Bioinformatic validation was also performed. Using another dataset, the KIF11 expression value was significantly different in the expression samples derived from SCLC cell lines. Another factor that may have an impact on the KIF11 transcription process is DNA methylation (Győrffy et al. 2016). After the exploration of the methylation array, one probe related to KIF11 showed a statistical difference, whereas the other did not. However, the significance level of methylation is far less than the difference in miRNA expression, indicating the susceptibility of the miRNA-mRNA pair in coordinating KIF11 expression (Fig. 10).

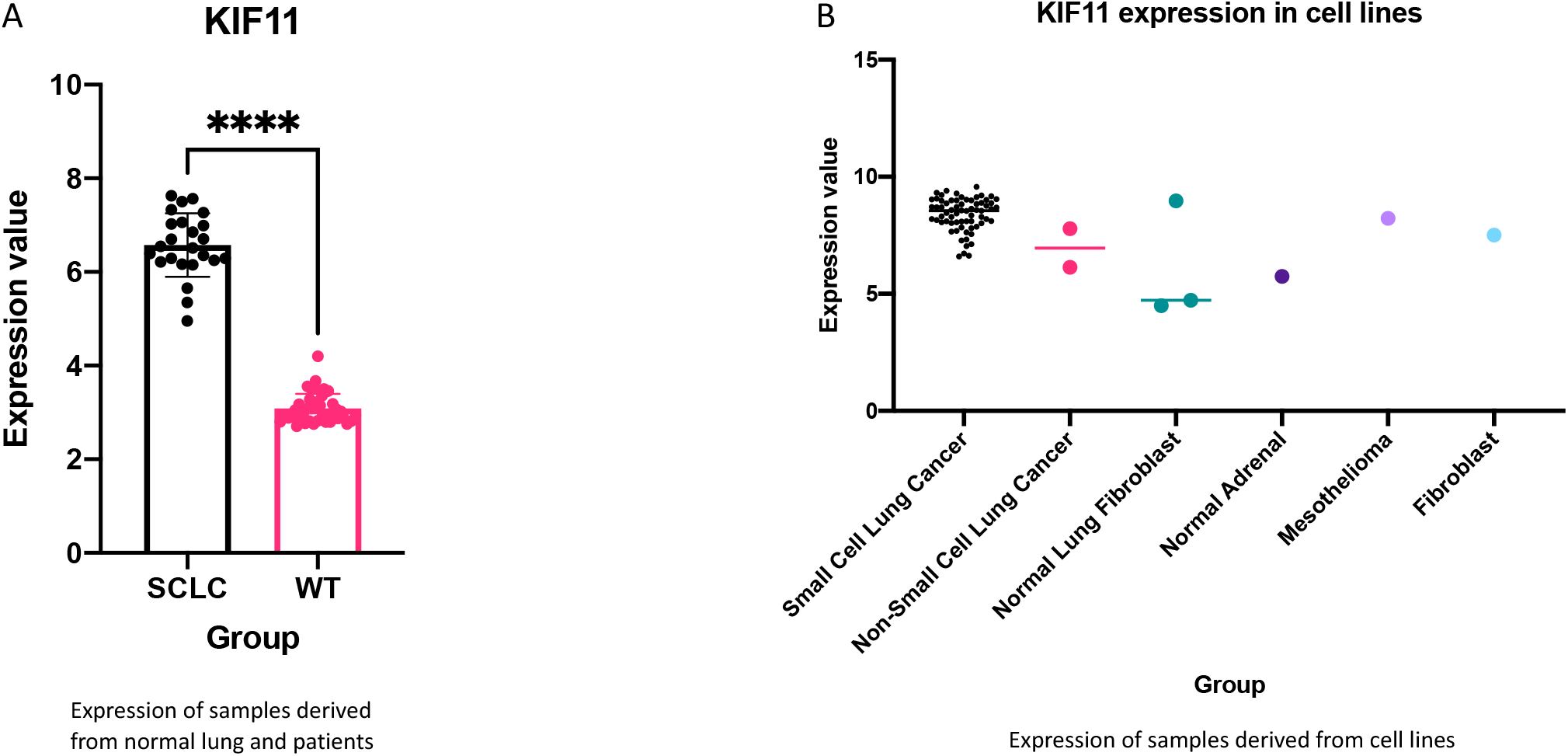

Because the unsupervised clusters based on ferroptosis genes and hub genes are nearly identical, it is reasonable to assume that the hub genes may have an impact on ferroptosis. The cell cycle may be a factor in the ferroptosis process. Moreover, the PPI between KIF11 and GPX4 is provided in the STRING database. The interaction was discovered by text-mining(Chevalier et al. 2012; Reemann et al. 2014; Rolland et al. 2009; Yang et al. 2016). As mentioned earlier, KIF11 plays an acceptable role and can be used as a therapeutic target. This makes the identification of this miRNA-mRNA pair more important for biomedical applications.

We did not provide any biological validation of miRNA-mRNA pairs, which is a limitation of this study. We identified miRNA-mRNA pairs using the miRTarbase database, which is a special database for identifying miRNA-mRNA targeting relationships supported by experimental evidence. The database is based on natural language processing technology (Hsu et al. 2011). Therefore, the interaction has already been proven by other studies.

However, future analysis should further experiment with the KIF11 function in SCLC, based on the aim of this study to provide a novel therapeutic approach and target for SCLC treatment. The cluster method provided in this analysis is also a type of unsupervised learning process. Based on the limitations of SCLC data, the results may have a type 2 error. However, the novel logical method is more than essential for learning and thus applicable to future analyses. The results would also be more accurate if more SCLC data are generated in the future.

## 5. Conclusion

In this *in silico* analysis, hub genes in SCLC are identified. Differentially expressed miRNAs are also shown. The important roles of the miR-30 family and KIF11 pair are discussed. KIF11 may also play a role in ferroptosis. To the best of our knowledge, the detected miRNA-mRNA pair provides a novel therapeutic approach for SCLC.

## Acknowledgments

Yuefeng would like to thank everyone in Songlab for discussion.

## Author Contributions

Yuefeng Wu conceived the idea and designed the experiments

## Data Availability

The data can be accessed in GEO database.

