## Supplementary material for "Identification and verification of the potential effect of therapeutic miRNA-mRNA pairs on ferroptosis in small cell lung cancer by bioinformatics analysis": Figure Legends

Figure legend：

Fig. 1 Hub genes and their protein-protein interactions.

Fig. 2. KEGG and GO enrichment results. The horizontal axis shows the enrichment ratio. The shape of the dots indicates the gene counts, while the color represents the significance of the enrichment results.

Fig. 3 Delta area curve of consensus clustering, indicating the relative change in area under the cumulative distribution function (CDF) curve for each category number k compared with k–1. The horizontal axis represents the category number k and the vertical axis represents the relative change in area under the CDF curve.

Fig. 4 Hub gene: Heatmap depicting consensus clustering solution (k = 3) for 14 hub genes in 24 samples (n = 24). Heatmap of hub gene expression in different subgroups: red represents high expression and blue represents low expression. Ferroptosis gene: Heatmap depicting the consensus clustering solution (k = 3) for 24 ferroptosis-related genes in 24 samples (n = 24).

Fig. 5 C)Venn map showing the crosstalk between corresponding unsupervised cluster groups. Here, different colors represent different groups. A, B, D）The ferroptosis related genes expressing value in sub groups. Asterisks represent levels of significance; *p < 0.05, **p < 0.01, ***p < 0.001.

Fig. 6 Volcano map showing the differentially expressed miRNAs

Fig. 7 KEGG and GO enrichment results. The horizontal axis indicates the p-value. The shape of the dots and colors show the gene counts.

Fig. 8 A) Venn map with the crosstalk between target genes and hub genes. B) indicating the miR-30 family expression difference between SCLC and normal cells.

Fig. 9 Methylation of KIF11 status between SCLC and normal cells.

Fig. 10 Comparison of KIF11 expression in patients with SCLC (A) and other cell lines (B).
